# Radicle growth regulation of root parasitic plants by auxin-related compounds

**DOI:** 10.1101/2024.05.03.592385

**Authors:** Kei Tsuzuki, Taiki Suzuki, Michio Kuruma, Kotaro Nishiyama, Ken-ichiro Hayashi, Shinya Hagihara, Yoshiya Seto

## Abstract

Root parasitic plants in the Orobancheceae, such as *Striga* and *Orobanche*, cause significant damage to crop production. The germination step of these root parasitic plants is induced by host-root-derived strigolactones (SLs). After germination, the radicles elongate toward the host and invade the host root. We have previously discovered that a simple amino acid, tryptophan (Trp), as well as its metabolite, the plant hormone indole-3-acetic acid (IAA), can inhibit radicle elongation of *Orobanche minor*. These results suggest that auxin plays a crucial role in the radicle elongation step in root parasitic plants. In this report, we used various auxin chemical probes to dissect the auxin function in the radicle growth of *O. minor* and *Striga hermonthica*. We found that synthetic auxins inhibited radicle elongation. In addition, auxin receptor antagonist, auxinole, rescued the inhibition of radicle growth by exogenous IAA. Moreover, a polar transport inhibitor of auxin, *N*-1-naphthylphthalamic acid (NPA), affected radicle tropism. We also proved that exogenously applied Trp is converted into IAA in *O. minor* seeds, and auxinole partly rescued this radicle elongation. Our data demonstrate a pivotal role of auxin in radicle growth. Thus, manipulation of auxin function in root parasitic plants should offer a useful approach to combat these parasites.

## Introduction

Root parasitic plants, such as *Striga* and *Orobanche* species, cause critical damage to crop production, particularly in sub-Saharan Africa. They germinate only in the presence of their host plants nearby by sensing strigolactones (SLs) that are exudated from the host roots (Xie et al. 2010). The parasitic plant seeds can stay dormant for decades, but once the host plant occurs nearby, they germinate by sensing SLs. This unique germination mechanism is thought to be a survival strategy for the parasitic plants, which are not able to survive without attaching to the host. On the basis of this germination mechanism, suicidal germination induction has been proposed as an effective way to eliminate parasitic plant seeds from infested fields. Toward the practical application of this method, many synthetic analogs of SLs have been synthesized (Tsuchiya 2018). Recently, SL receptors in root parasitic plants, such as *Striga hermonthica*, *Phelipanche ramosa*, and *Orobanche minor*, have been characterized (Conn et al. 2015; de Saint Germain et al. 2021; Takei et al. 2023; Toh et al. 2015; Tsuchiya et al. 2015), and this has facilitated the identification of novel chemicals that target these receptors (Uraguchi et al. 2018; Wang et al. 2022). Such synthetic agonists can be used as suicidal germination inducers that promote germination in the absence of the host plants. Since they cannot survive without a host, germinated seeds die within 4 to 5 days. Although this method is effective, because of the difficulty of mass production at low cost, it has not been put into practical use.

After root parasitic plant seeds germinate in an SL-dependent manner, their radicle elongates toward the host root. A recent report demonstrated that SLs play a critical role not only as germination stimulants but also as chemoattractants that induce a host-tropic response in the elongating radicle (Ogawa et al. 2022). Considering its importance, the radicle elongation step might be a promising target to inhibit parasitism. We have previously found that a simple amino acid, tryptophan (Trp), has inhibitory activity on the radicle elongation of *O. minor*. Because Trp is a precursor of a natural auxin, indole-3-acetic acid (IAA), we also investigated the activity of IAA and showed that IAA inhibited the radicle growth of *O. minor* (Kuruma et al. 2021). However, it remained unclear whether IAA inhibited the radicle growth by modulating auxin signaling pathway. Moreover, if radicle elongation depends on auxin signaling, it would be interesting to know how it regulates the radicle growth. To address these questions, in the current work we examined the effects of chemical probes on auxin response during the radicle growth of two root parasitic plants, *O. minor* and *S. hermonthica*. Our results demonstrate that exogenous auxin can inhibit radicle elongation of the parasitic plants and that endogenous auxin might have an important function in regulating the tropic response of the radicle.

## Results

### Effects of the synthetic auxin analogs, 2,4-D and NAA

As outlined above, our previous results demonstrated that IAA inhibited the post-germination radicle growth of *O. minor*, when co-treated with a synthetic SL analog, *rac*-GR24 (Kuruma et al. 2021). To examine whether IAA inhibits the radicle growth by activating auxin signaling pathway, we tested the activity of commercially available synthetic auxins, 2,4-dichlorophenoxyacetic acid (2,4-D) and 1-naphthaleneacetic acid (NAA). The germination of *O. minor* was induced by *rac*-GR24, in the presence of 2,4-D or NAA, and we found that both these two synthetic auxins inhibited post-germination radicle elongation of *O. minor* in a concentration-dependent manner, as did IAA (Fig. 1A). The same effect was observed when *S. hermonthica* seeds were treated with 2,4-D or NAA (Fig. 1B). These results demonstrated that radicle growth would be modulated by auxin signaling activation, and thus excess auxin treatment had a negative effect on the radicle elongation of both *O. minor* and *S. hermonthica*.

**Fig. 1.**
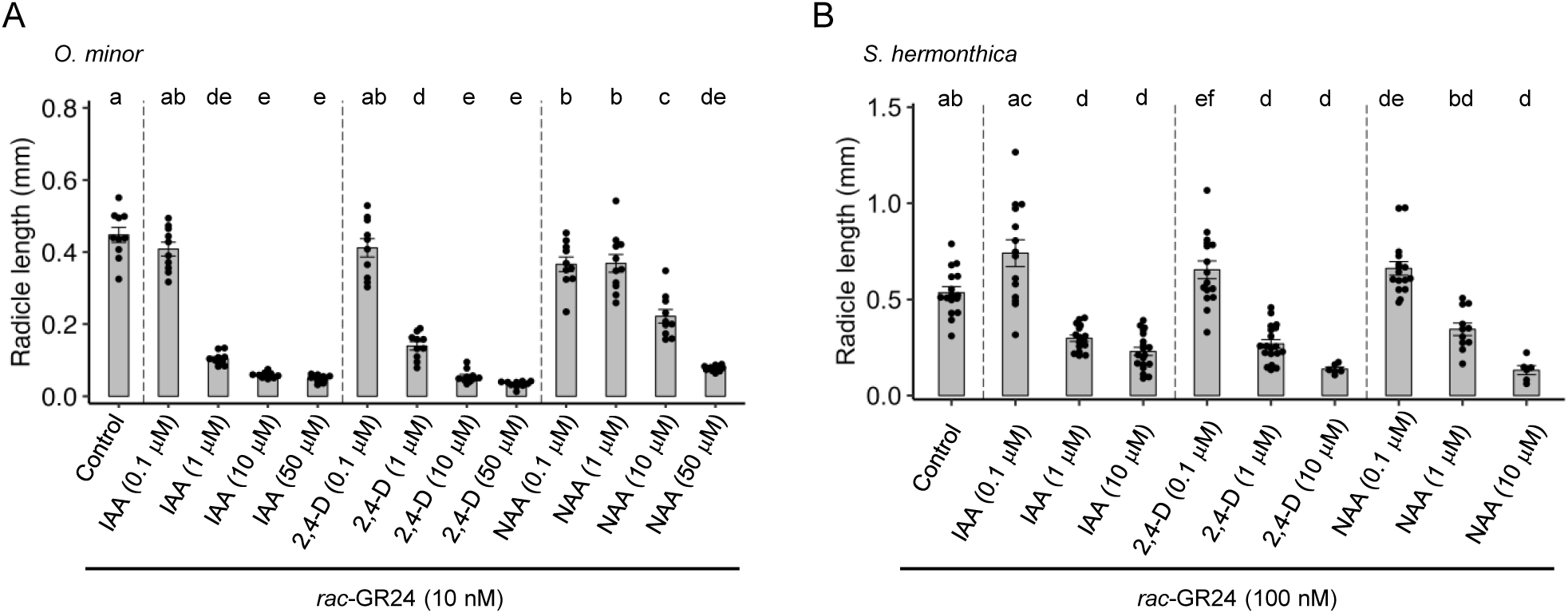
Effects of auxin agonists on radicle growth of germinated *O. minor* seeds (**A**) or *S. hermonthica* seeds (**B**). Data are the means ± SE (*n* = 10 in (A) and *n* = 6-19 in (B)). 10 or 100 nM *rac*-GR24 solution contained 0.1% acetone was used as control. Different letters indicate significant differences at *P* < 0.05 with Tukey multiple comparison test. Dots show the exact data for individual samples.

### Effects of the IAA transport inhibitor, NPA

Our results suggested that exogenous auxin can inhibit the radicle elongation of root parasitic plants, and that perturbation of auxin responses in parasitic plants might be a promising approach to control their radicle growth. Therefore, we next evaluated chemicals that affect endogenous auxin function. It is well known that polar transport is important for auxin function, and *N*-1-naphthylphthalamic acid (NPA) is auxin transport inhibitor that is widely used for diverse plants. When *O. minor* seeds were co-treated with NPA plus *rac*-GR24, we did not observe the inhibitory effects on radicle elongation below 1 μM (Fig. 2A). At higher concentrations, NPA moderately inhibited radicle elongation (Fig. 2A). When *O. minor* seeds were treated with SL alone, the germinated radicles elongated, and often bent in a random direction. However, we noticed that, even with treatment at 1 μM NPA, the radicles grew straighter than in the untreated control. We measured the angle of the elongated radicles, and found the angle of the radicle elongating direction was indeed smaller in the presence of NPA (Fig. 2B, Fig S1). These results suggested that the endogenous auxin transport played a critical role in regulating the radicle tropic response.

**Fig. 2.**
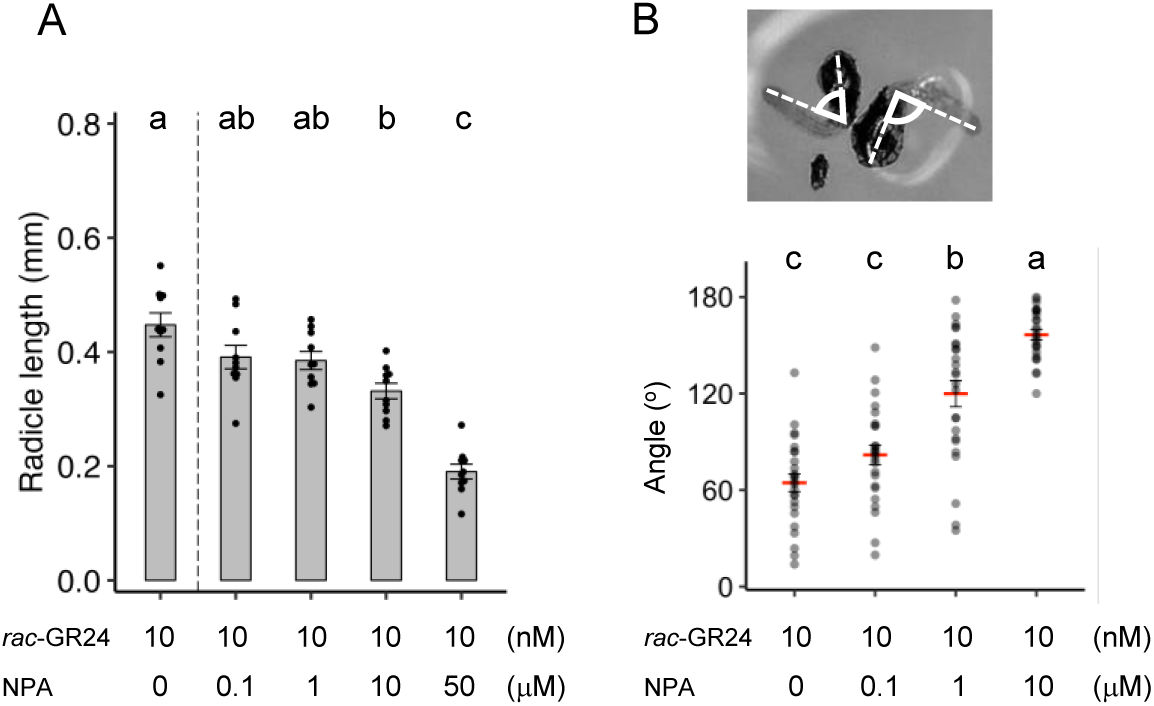
Effects of the auxin transport inhibitor on radicle growth or angle of germinated *O. minor* seeds. (A) Radicle growth inhibitory activity of NPA. Data are the means ± SE (*n* = 10). (B) The radicle angle of germinated *O. minor* seeds in the presence of NPA. Data are the means ± SE (*n* = 25). Different letters indicate significant differences at *P* < 0.05 with Tukey multiple comparison test. Dots show the exact data for individual samples.

### Effects of the *cis* isomer of cinnamic acid

In addition to NPA, the *cis* isomer of cinnamic acid (CA) was previously reported to affect polar auxin transport (Steenackers et al. 2017). CA normally exists as the *trans* form but UV irradiation stimulates its isomerization to the *cis* isomer. *cis*-CA occurs in plants as an endogenous compound, yet how it is produced *in planta* and its endogenous function have not been fully elucidated. A previous report demonstrated that *cis*-CA treatment at low concentrations, such as 1 μM, promoted the growth of *Nicotiana benthamiana* or *Arabidopsis* by manipulating endogenous auxin function (Steenackers et al. 2019). At higher concentrations (above 10 μM), *cis*-CA led to growth inhibition in both species. Thus, we expected that *cis*-CA treatment of root parasitic plants would have some effects on radicle growth. We prepared *cis*-CA by UV irradiation of commercially available *trans*-CA as we have previously reported (Suzuki et al. 2022), and tested its activities on *O. minor* radicle elongation. We found that *cis*-CA inhibited radicle elongation of *O. minor* in a dose-dependent manner, but *trans*-CA did not (Fig. 3A). The similar response to *cis*-CA was also observed in *S. hermonthica* (Fig. 3B).

**Fig. 3.**
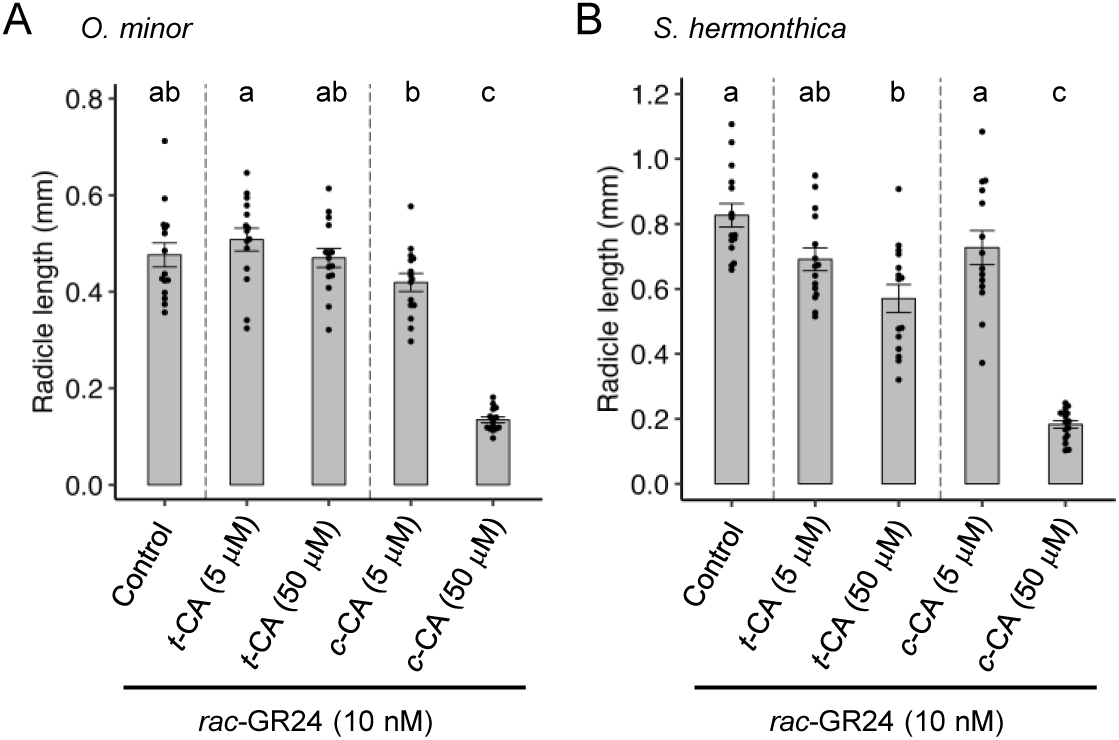
Effects of the *trans*- or *cis*-cinnamic acid on radicle growth of germinated seeds of *O. minor* (**A**) or *S. hermonthica* (**B**). Data are the means ± SE (*n* = 14-16). 10 nM *rac*-GR24 solution contained 0.1% acetone was used as control. Different letters indicate significant differences at *P* < 0.05 with Tukey multiple comparison test. Dots show the exact data for individual samples.

### Effects of the IAA antagonist, auxinole

Next, to address physiological roles of endogenous auxin during the radicle elongation, we examined the effects of auxin receptor antagonist, auxinole during the *O. minor* germination (Hayashi et al. 2012). We found that auxinole did not affect radicle elongation (Fig. 4A). It was possible that auxinole was not recognized by auxin TIR1/AFB receptors in *O. minor*. To examine this possibility, we co-treated with auxinole plus IAA. We found that auxinole rescued the IAA-inhibited radicle growth in *O. minor* (Fig. 4B, Fig S2). These results suggested that endogenous IAA level would be lower than the effective concentration required for the radicle growth suppression.

**Fig. 4.**
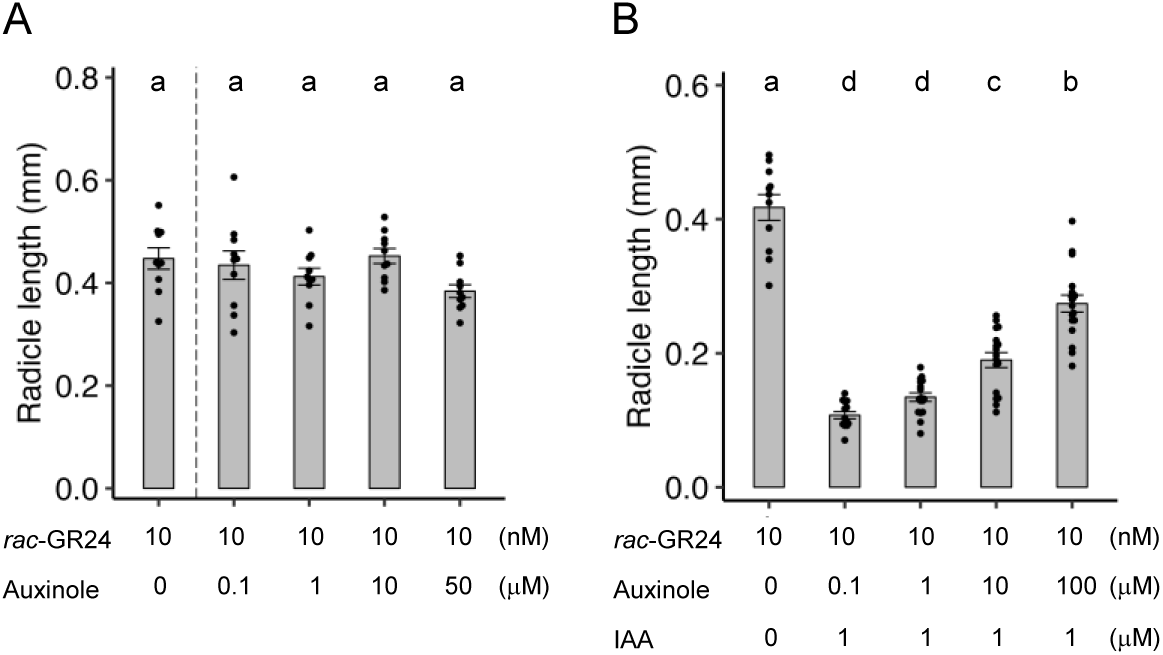
Effects of auxin antagonist on radicle growth of germinated *O. minor* seeds. (**A**) Radicle growth inhibitory activity of auxinole. Data are the means ± SE (*n* = 10). (B) The antagonistic activity of auxinole to exogenously applied IAA. Data are the means ± SE (*n* = 11-18). Different letters indicate significant differences at *P* < 0.05 with Tukey multiple comparison test. Dots show the exact data for individual samples.

### Effects of the C5-substituted IAA analogs

The above-mentioned results strongly suggested that artificial regulation of the endogenous auxin function in root parasitic plants might be a promising tool to control their radicle growth. Exogenous treatment with IAA, 2,4-D, or NAA was effective for inhibiting the radicle growth of root parasitic plants. However, these auxins are active in both the root parasitic plants and the host plants. If considering the use of auxin-related compounds as agrochemicals, we preferably need selective chemicals that function only in the root parasitic plants. Aiming to identify such chemicals, we focused on a series of synthetic auxin analogs that have a substitution at the C-5 position. Previous reports demonstrated that introducing a substitution at the C-5 position of IAA attenuated its auxin activity (Uchida et al. 2018; Yamada et al. 2018). Compared with IAA, these C-5 substituted IAA derivatives showed weaker activity in inducing the receptor complex formation between the receptor, TIR1, and its repressor protein (Uchida et al. 2018). Interestingly, mutant TIR1 proteins, in which a specific Phe residue at the ligand binding pocket was replaced by a small amino acid such as Ala or Gly, were able to bind with the C-5 substituted analogs. *Arabidopsis* WT plants were resistant to the C-5 substituted IAA analogs; however, transgenic *Arabidopsis* plants expressing the mutant TIR1 were able to respond to the IAA derivatives. Therefore, if the C-5 substituted IAA analogs retained activity toward root parasitic plants, such chemicals might be good candidates as agrochemicals that would specifically control the growth of root parasitic plants without affecting the host plant growth.

Among C-5 substituted auxin analogs, cvx-IAA and adamantyl-IAA (ada-IAA) are commercially available, and therefore we tested the activity of these compounds on the radicle growth of *O. minor*. Interestingly, both IAA analogs inhibited radicle elongation at moderately low concentrations (Fig. 5A). In particular, ada-IAA showed stronger activity than cvx-IAA, and its activity was almost as same as that of IAA. In addition, we found that these IAA analogs inhibited radicle elongation of *S. hermonthica* (Fig. S3). We also tested the effects of these compounds on a non-parasitic plant, *Arabidopsis*. Both IAA analogs showed much weaker activity in inhibition of primary root elongation in *Arabidopsis*, compared with IAA (Fig. 5B, Fig. S4). Although treatment with ada-IAA did not inhibit primary root growth in *Arabidopsis*, we found that the root growth direction was affected in the presence of ada-IAA, suggesting that ada-IAA might affect the gravitropic response selectively without affecting the extent of root growth (Fig. S4). Although, cvx-IAA did not affect the gravitropism at physiological concentrations (<0.3 μM), we found that at high concentration it affects the gravitropic response as was previously suggested (Fig. S4) (Uchida et al. 2018). Such effects of C-5 substituted auxin analogs were reported using C-5 alkoxy analogs, and these analogs were found to inhibit auxin transports selectively (Tsuda et al. 2011). Therefore, we expect that ada-IAA or cvx-IAA also selectively inhibits auxin transporters.

**Fig. 5.**
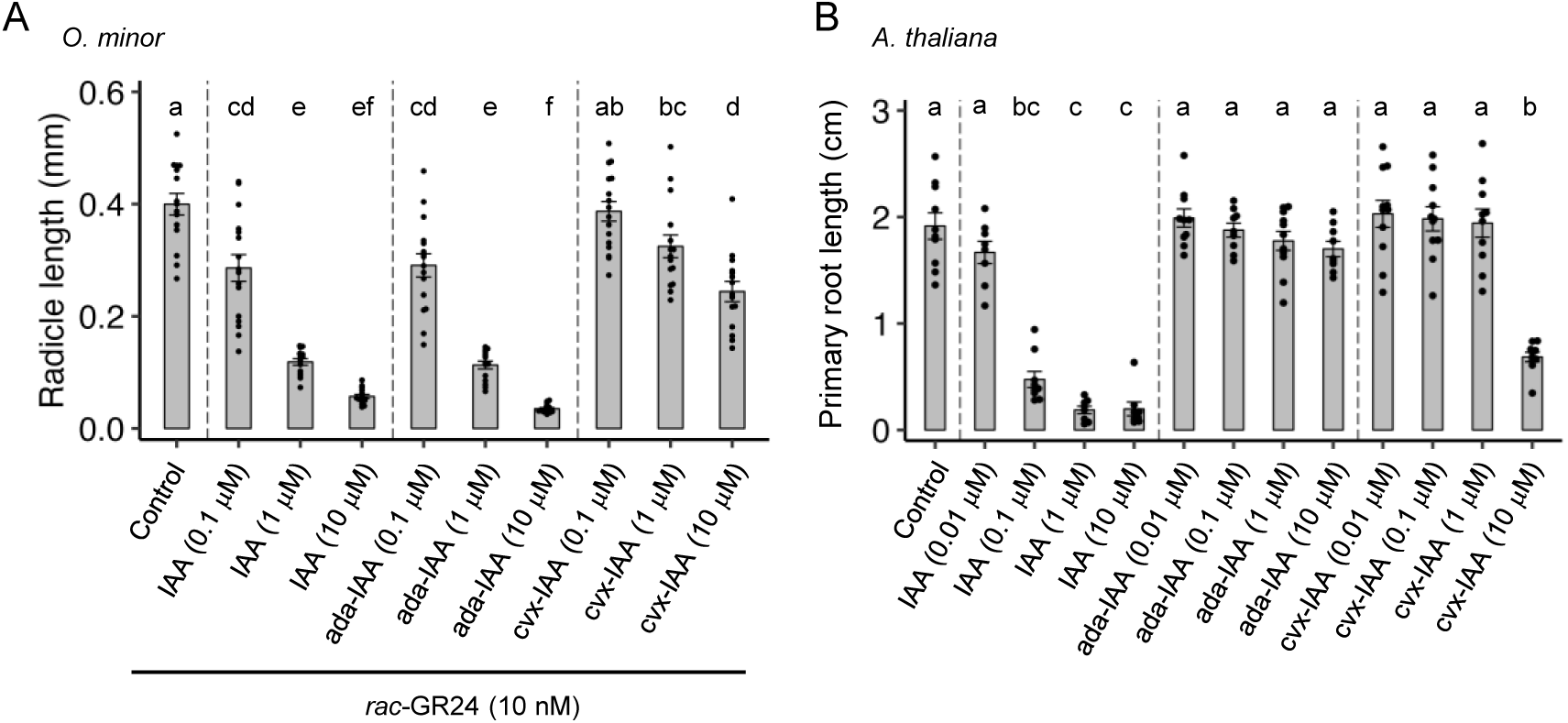
Effects of synthetic auxin analogs on radicle growth of germinated *O. minor* seeds or primary root growth of *A. thaliana* seedling. (A) Radicle growth inhibitory activity of synthetic auxin analogs for *O. minor* seeds. 10 nM *rac*-GR24 solution contained 0.1% acetone was used as control. Data are the means ± SE (*n* = 15-16). (B) Inhibitory activity of synthetic auxin analogs on primary root growth of *A. thaliana* seedlings. ½ MS agar plate contained 0.1% acetone was used as control. Data are the means ± SE (*n* = 8-11). Different letters indicate significant differences at *P* < 0.05 with Tukey multiple comparison test. Dots show the exact data for individual samples.

### Conversion of tryptophan into indole-3-acetic acid in *Orobanche minor*

Our results clearly demonstrated that auxin treatment can inhibit the radicle growth of root parasitic plants. We had previously identified a simple amino acid, Trp, as a radicle growth inhibitor. Because Trp is a biosynthetic precursor of IAA in plants, we expected that excess Trp would be converted into IAA *in planta*, and then inhibits radicle growth. To examine this hypothesis, we performed a feeding experiment using a stable isotope-labeled Trp (*d*_5_-Trp). Conditioned *O. minor* seeds were cultured with GR24 and *d*_5_-Trp and incubated for 24 h. An acetone extract of the seeds was then analyzed by LC–MS/MS. We successfully detected a peak whose molecular weight was identical to that of *d*_5_-IAA (Fig. 6). Moreover, the retention time and the MS/MS spectrum supported the identity of the peak as *d*_5_-IAA (Fig. 6). These results strongly suggested that exogenously applied Trp was at least partly converted into IAA in the *O. minor* seeds, supporting our hypothesis.

**Fig. 6.**
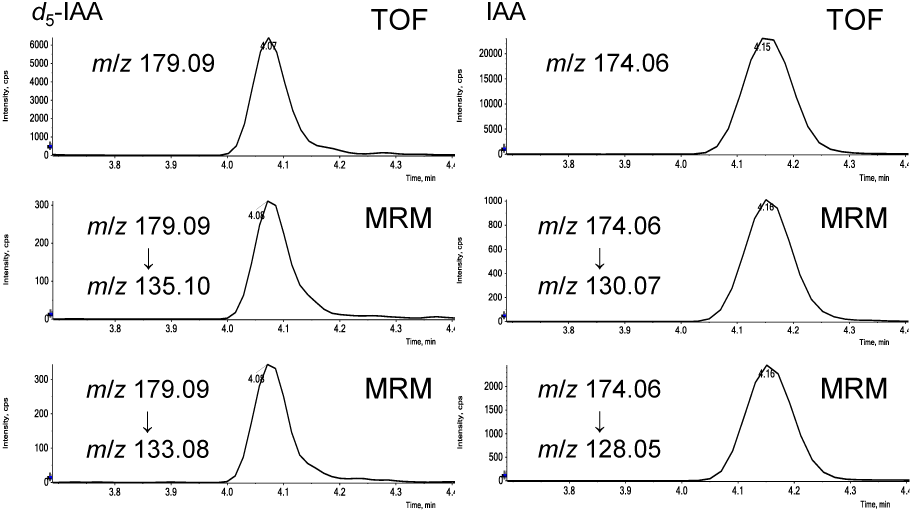
LC-MS/MS analysis of *d*_5_-IAA from acetone extracts of *O. minor* seeds co-treated with GR24 and *d*_5_-Trp for 5 days.

### Effects of IAA biosynthetic inhibitors

Exogenously applied Trp was shown to be converted into IAA. Therefore, it would be possible that the radicle growth inhibition by Trp was caused by IAA which was converted from the applied Trp. If this were the case, co-treatment with Trp plus inhibitors of IAA biosynthesis might be able to rescue the radicle growth inhibition. To investigate this hypothesis, we first tested effect of the known IAA biosynthetic inhibitors, yucasin DF (YDF) and kynurenine (KYN) to *O. minor* (He et al. 2011; Tsugafune et al. 2017). However, the endogenous IAA level was not decreased after co-treatment of YDF and KYN (Fig. 7A). We also found that these IAA biosynthetic inhibitors did not affect radicle growth of *O. minor* (Fig. 7B). These IAA biosynthetic inhibitors were originally discovered using IAA biosynthetic enzymes in *Arabidopsis*. Therefore, our results indicate that these inhibitors do not affect the IAA biosynthetic enzymes in *O. minor*.

**Fig. 7.**
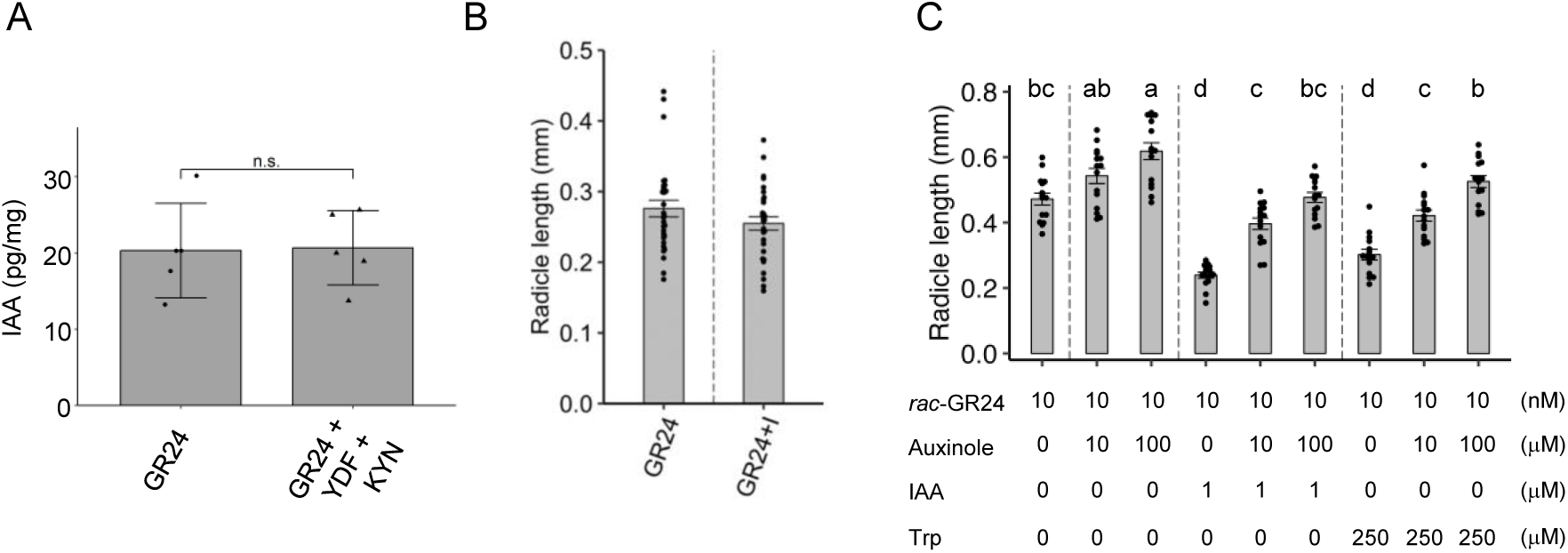
Effects of auxin biosynthesis inhibitors or antagonist. (**A**) Effects of IAA biosynthetic inhibitors, yuccasin DF (YDF) and kynurenine (KYN), on the radicle elongation in *O. minor*. GR24+I means GR24 and two inhibitors. (**B**) Quantitative analysis of endogenous level of IAA from acetone extract of *O. minor* seeds after co-treatment of GR24 and IAA biosynthesis inhibitors. GR24: 100 nM GR24, GR24+YDF+KYN: 100 nM GR24, 50 μM YDF, 10 μM KYN. Data are means ± SD (n=5). T-test: n.s. > 0.05. (**C**) Effects of auxinole on radicle growth of *O. minor* seeds in the presence of exogenously applied IAA or Trp. Data are the means ± SE (*n* = 15). Different letters indicate significant differences at *P* < 0.05 with Tukey multiple comparison test. Dots show the exact data for individual samples.

As mentioned above, co-treatment with IAA together with auxinole rescued radicle growth inhibition. Thus, we next investigated the effect of auxinole on the radicle growth inhibition induced by Trp. We found a partial rescue of radicle growth inhibition, compared with treatment with Trp alone (Fig. 7C). These results, together with demonstrating Trp conversion into IAA, suggested that the radicle growth inhibition caused by Trp treatment is, at least in part, caused by IAA converted from administered Trp.

### Synthesis and biological activities of auxin-derived strigolactone-type hybrid compounds

As mentioned above, SLs induce the germination of root parasitic plants. Because SL synthetic analogs might be used as suicidal germination inducers for root parasitic plants, numerous SL analogs have been reported. Most of these chemicals simply induce the germination of root parasitic plants by mimicking SL activity. In contrast, we have previously reported a hybrid-type SL analog, IAA–SL, in which IAA is attached to methylbutenolide (commonly referred to as D-ring), which is the typical substructure in SL molecules. Recent reports have demonstrated that this part of the molecule is essential for SL activity, while the ABC-ring part of the structure has a low functional requirement. The hybrid compound, IAA–SL, retains the germination-inducing activity toward *O. minor* (Hylova et al. 2019; Kuruma et al. 2021). Moreover, the IAA part can inhibit radicle elongation. Thus, IAA–SL has two activities that induce germination and inhibit subsequent radical growth of *O. minor* (Kuruma et al. 2021). The results described earlier suggested that 2,4-D, NAA, and ada-IAA would also be good candidates for synthesizing such hybrid compounds. Accordingly, we chemically synthesized three hybrid compounds, 2,4-D–SL, NAA–SL, and ada-IAA–SL (Fig. 8A). All of these chemicals induced the germination of *O. minor* seeds, yet the activity of 2,4-D-SL or ada-IAA–SL was much weaker compared with 2,4-D–SL and NAA–SL (Fig. 8A). In addition, these chemicals showed radicle growth-inhibiting activity at relatively high concentrations (Fig. 8B). Compared with IAA–SL, these newly synthesized compounds showed weaker activity in terms of both germination induction and radicle growth inhibition.

**Fig. 8.**
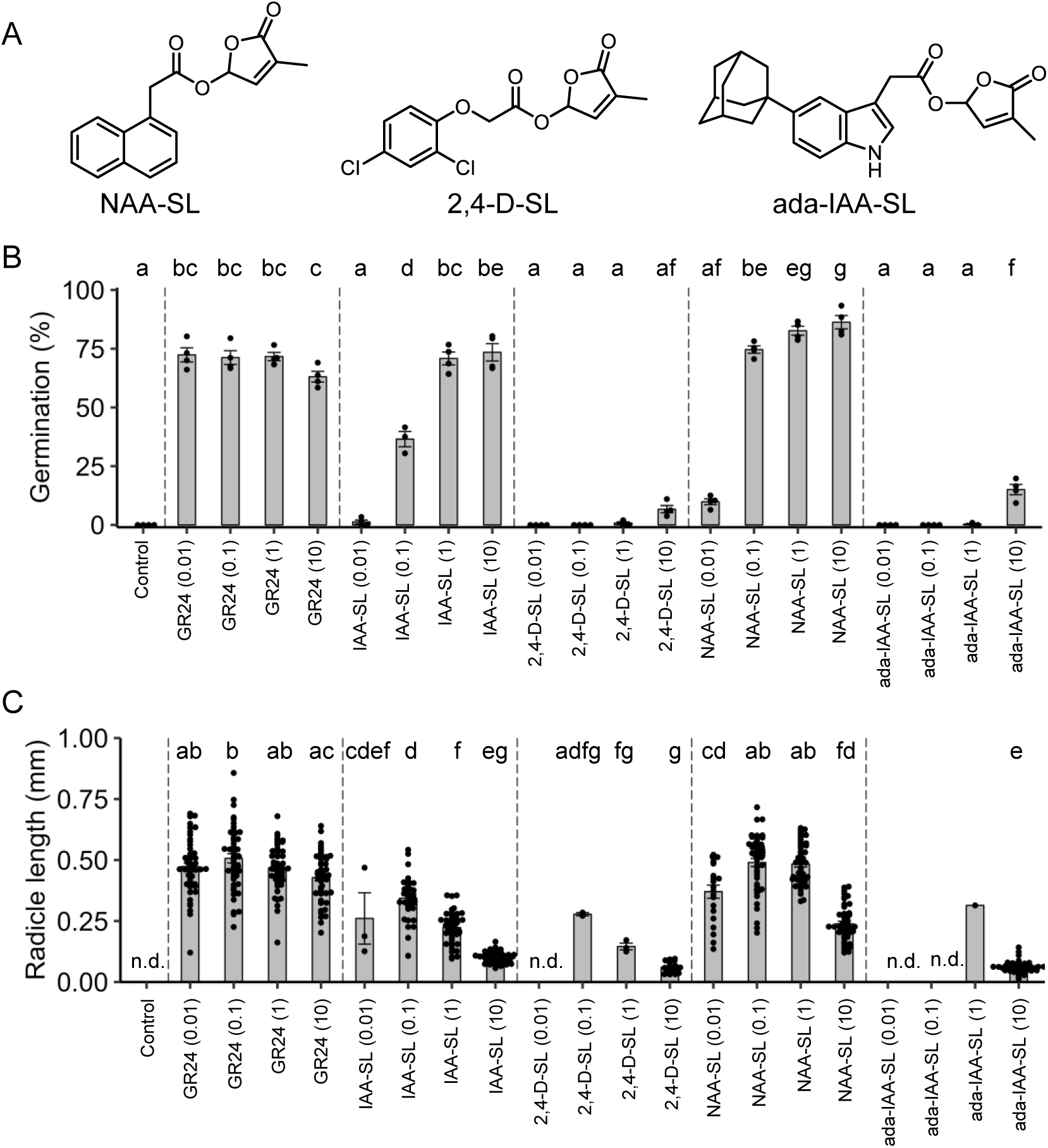
Effect of synthetic auxin-SL hybrid compounds on the germination and radicle elongation of *O. minor*. (**A**) Chemical structures of the synthesized hybrid compounds. (**B**) Germination-inducing activity of the auxin-SL hybrid compounds toward *O. minor* seeds. Data are the means ± SE (*n* = 3-4). Different letters indicate significant differences at *P* < 0.05 with Tukey multiple comparison test. Dots show the exact data for individual samples. Control means that 0.1% acetone was administered. The values in parentheses indicate the concentration.(**C**) Effect of auxin-SL hybrid compounds on post-radicle elongation on *O. minor*. Data are the means ± SE (*n* = 1-48). Different letters indicate significant differences at *P* < 0.05 with Tukey multiple comparison test. Dots show the exact data for individual samples. Control means that 0.1% acetone was administered and n.d. indicates “not detected”, because no seed germinated. The values in parentheses indicate the concentration.

## Discussion

We previously reported that Trp or IAA inhibited the radicle elongation of *O. minor*. In this report, we demonstrate that not only IAA but also 2,4-D and NAA had the same activity as IAA, suggesting that auxin inhibited post-germination radicle growth of *O. minor* and *S. hermonthica*. In addition, labeled *d*_5_-Trp was converted into *d*_5_-IAA in *O. minor* seeds. Moreover, the recovery of radicle elongation inhibition by an auxin antagonist, auxinole, was observed not only when co-treated with IAA, but also when co-treated with Trp. Taken together, our data strongly suggested that the radicle inhibition effect of Trp treatment was, at least in part, caused by IAA converted from Trp.

We also found that polar auxin transport inhibitors affected the radicle growth of *O. minor* and *S. hermonthica*. In particular, NPA did not inhibit radicle elongation at lower concentrations. However, even at such concentrations, NPA inhibited the radicle tropism of *O. minor*. Moreover, *cis*-CA, but not *trans*-CA, also showed radicle elongation inhibiting activity. A previous report suggested that the root parasitic plant radicle showed a tropic response to SL, and that polar auxin transport would be important for this response (Ogawa et al. 2022). Our results support this working model, and therefore indicate that polar auxin transport machinaries would be a promising target to control the radicle elongation step of colonization by root parasitic plants.

Manipulation of auxin regulated pathways in root parasitic plants might be a useful approach to combat these parasites. However, it might be difficult to control root parasitic plants selectively without affecting the host growth. In this regard, C-5 substituted IAA analogs, such as ada-IAA, are good candidates to overcome this problem. Among them, ada-IAA was previously reported as a potent ligand for a mutated TIR1 receptor (Uchida et al. 2018; Yamada et al. 2018). We found that this chemical inhibited radicle elongation of *O. minor* and *S. hermonthica*, while it did not inhibit the primary root growth of *Arabidopsis*. Therefore, ada-IAA is a good potential lead compound to develop an agrochemical that might selectively target root parasitic plants. We found that root gravitropism was weakly affected by ada-IAA, suggesting that ada-IAA retains partial activity in *Arabidopsis* as was the case with C-5 alkoxy auxin analogs (Tsuda et al. 2011). Therefore, it might be better to explore more specific chemicals that do not affect *Arabidopsis* root growth, but inhibit the radicle growth of root parasitic plants.

In conclusion, using various auxin chemical probes, we demonstrated here that auxin plays a crucial role in radicle growth regulation in root parasitic plants. Therefore, our results confirm that artificial control of auxin function is a worthwhile target for the development of new agrochemicals against root parasitic plants.

## Materials and Methods

### Plant materials

We used *Arabidopsis* ecotype Col-0. *Orobanche minor* seeds were harvested from Nagasawa water purification plant, Kawasaki, Japan. *Striga hermonthica* seeds were kindly provided from Dr. Steven Runo.

### Chemicals

Ada-IAA, cvx-IAA, and kynurenine are commercially available. Preparation of yucasin DF was previously reported (Tsugafune et al. 2017).

### Chemical Synthesis

*cis*-CA, 5-bromo-3-methyl-2(5H)-franone and IAA-SL were prepared according to our previous reports (Suzuki *et al*., 2022; Kuruma *et al*., 2021).

2,4-D–SL; K_2_CO_3_ (1.0 mmol) was added to the solution of 2,4-D (0.50 mmol) in *N*-methyl-2-pyrrolidone (5 mL). 5-bromo-3-methyl-2(5H)-franone (0.98 mmol) diluted in *N*-methyl-2-pyrrolidone (5 mL) was added to the mixture. After stirring for 16 h at room temperature, the reaction was quenched by adding 1 N HCl, and the solution was diluted to 50 mL with water. The solution was extracted with EtOAc (3×50 mL). The organic layer was washed with distilled water (2×150 mL), dried over Na_2_SO_4_, and concentrated *in vacuo*. The crude sample was purified by silica gel column chromatography (*n*-hexane/EtOAc:7/3) to afford 2,4-D–SL (84.8 mg, 0.27 mmol, 53%). ^1^H-NMR (300 MHz, CDCl_3_) *δ* 7.40 (d, *J* = 2.1 Hz, 1H), 7.19 (dd, *J* = 8.7 Hz, *J* = 2.7 Hz, 2H), 6.97-6.95 (m, 1H), 6.94-6.92 (m, 1H), 6.82 (d, *J* = 9.0 Hz, 1H), 4.76 (s, 3H), 2.01-2.00 (m, 3H); ^13^C-NMR (75 MHz, CDCl_3_), *δ* 10.69, 66.13, 92.70, 115.35, 124.50, 127.73, 130.46, 135.02, 141.30, 152.08, 166.70, 170.64; HRMS [ESI+ (*m*/*z*)] calculated for (C_13_H_10_Cl_2_O_5_ +H)^+^ 316.9978, found 316.9990.

NAA–SL; K_2_CO_3_ (1.0 mmol) was added to the solution of NAA (0.50 mmol) in *N*-methyl-2-pyrrolidone (5 mL). 5-Bromo-3-methyl-2(5H)-franone (0.98 mmol) diluted in *N*-methyl-2-pyrrolidone (5 mL) was added to the mixture. After stirring for 16 h at room temperature, the reaction was quenched by adding 1 N HCl, and the solution was diluted to 50 mL with water. The solution was extracted with EtOAc (3×50 mL). The organic layer was washed with distilled water (2×150 mL), dried over Na_2_SO_4_, and concentrated *in vacuo*. The crude sample was purified by silica gel column chromatography (*n*-hexane/EtOAc:7/3) to afford NAA–SL (41.9 mg, 0.15 mmol, 30%). ^1^H-NMR (300 MHz, CDCl_3_) *δ* 7.95-7.81 (m, 3H), 7.59-7.40 (m, 4H), 6.88-6.87 (m, 1H), 6.84-6.83(m, 1H), 4.14 (s, 2H), 1.97-1.95 (m, 3H); ^13^C-NMR (75 MHz, CDCl_3_), *δ* 10.55, 38.54, 92.64, 123.40, 125.42, 125.89, 126.55, 128.17, 128.45, 128.78, 129.01, 131.85, 133.76, 134.39, 169.73, 171.00; HRMS [ESI+ (*m*/*z*)] calculated for (C_17_H_14_O_4_ +H)^+^ 283.0965, found 283.0978.

5-Adamantyl-IAA–SL (ada-IAA-SL); K_2_CO_3_ (69 mg, 0.50 mmol) was added to the solution of ada-IAA (79 mg, 0.26 mmol) and 5-bromo-3-methyl-2(5H)-franone (91 mg, 0.53 mmol) in *N*-methyl-2-pyrrolidone (5 mL). After stirring for 14 h at room temperature, the reaction was quenched by adding 1 N HCl, and the solution was diluted to 25 mL with water. The solution was extracted with EtOAc (3×25 mL). The organic layer was washed with distilled water (3×75 mL), dried over Na_2_SO_4_, and concentrated *in vacuo*. The crude extract was purified by flash chromatography on a Biotage One instrument (SNAP ultra-column or Sfär D column, 7–70% *n*-hexane/EtOAc over 10 column volumes) to provide ada-IAA–SL (51 mg, 0.13 mmol, 49%), as a solid. ^1^H-NMR (500 MHz, CDCl_3_) *δ* 8.02 (s, 1H), 7.53 (s, 1H), 7.33-7.29 (m, 2H), 7.16-7.15 (m, 1H), 6.91-6.89 (m, 1H), 6.87-6.85 (m, 1H), 3.86 (s, 2H), 2.12 (s, 3H), 1.99-1.96 (m, 9H), 1.83-1.77 (m, 6H); ^13^C-NMR (125 MHz, CDCl_3_), *δ* 10.60, 29.10, 31.15, 36.09, 36.88, 43.82, 92.64, 98.27, 107.07, 114.17, 120.15, 123.34, 126.78, 134.27, 134.32, 142.11, 143.26, 170.16, 171.12; HRMS [ESI+ (*m*/*z*)] calculated for (C_25_H_27_NO_4_ +H)^+^ 406.2013, found 406.2016.

### Root phenotypic analysis of *Arabidopsis*

Sterilized seeds of the *Arabidopsis* Wild type Col-0 were soaked into sterilized water at 4°C for 2 days under dark condition. The seeds were put on the plate containing 1% (w/v) agar-solidified 0.5× Murashige and Skoog (MS) medium, 1% sucrose. Test compounds were dissolved in the medium at 1 mM by 10,000 times dilution from each acetone stock (final concentration of acetone was 0.01% (v/v)). These seedlings were cultured horizontally at 22°C for 9 days under LED light (105 mmol/m^2^/s) with a 16 h light / 8 h dark photoperiod. After the cultivation, pictures of the plants were taken, and the primary root length was measured by using ImageJ software.

### Germination assay

*Orobanche minor* or *Striga hermonthica* seeds were sonicated in 70% EtOH for 1 min. After removing 70% EtOH, 1% sodium hypochlorite solution containing 0.2% Tween-20 was added to the seeds, and sonicated for 4 min. To remove the sodium hypochlorite solution, the seeds were washed with a lot of sterilized water. The seeds were resuspended in 0.1% agar solution and loaded onto 5 mm glass fiber filter disks (20-100 seeds/disk) on paper filter. *O. minor* seeds were conditioned at 23°C for 14-15 days. S*. hermonthica* seeds were conditioned at 30°C for 7 days. After the conditioning, disks were transferred to 96-well plates. To prepare the chemical solution containing the auxin related compounds, each acetone stock solution of tested chemical was added to the plastic tube and acetone was evaporated by keeping the lid opened. After the dryness, chemical solutions containing *rac*-GR24 (final concentration of acetone was 0.1%) were added to each plastic tube. Plastic tubes were sonicated and vortexed to dissolve chemicals. A 30 mL aliquot of chemical solutions was added to the well. The 96-well plates were incubated for the same conditions as the conditioning. After 5 days from chemical treatment, pictures were taken under microscope and the number of total and germinated seeds of *O. minor* were counted. After 1 day or 2 days from chemical treatment, pictures were taken under microscope and the number of total and germinated seeds of *S. hermonthica* were counted. Radicle length and radicle angle were measured by using ImageJ software.

### Analysis of *d*_5_-IAA in *O. minor* seeds treated with *d*_5_-Trp

L-tryptophan-2,4,5,6,7-*d*_5_ (CDN isotope) was diluted to 1 mM with sterile water containing 0.1% acetone. 1 mM GR24 dissolved in acetone was diluted 1000-fold with sterile water to 1 nM. A sheet of filter paper (70 mm) was placed on a petri dish, followed by 1 mL each of 10 nM GR24 and 1 mM L-tryptophan-2,4,5,6,7-*d*_5_ to moisten the filter paper. Twenty conditioned *O. minor* seed discs were placed on it and excess water was removed. Petri dishes were closed with paraffin film, and incubated under dark at 23°C for 5 days. Then, the disks were placed on filter paper and washed with sterile water. 6 mL of acetone was added to the seeds on filter paper and extracted at 4°C overnight. The acetone solution was filtered, concentrated, and dissolved in 1 mL of 1% acetic acid/ water. This solution was loaded onto an HLB cartridge column (1 CC, 30 mg, Waters), washed with 1 mL of H_2_O containing 1% AcOH, and eluted with 2 mL of 80% MeCN containing 1% AcOH. The eluted fraction was concentrated again and dissolved in MeCN. The samples were subjected to LC-MS/MS and analyzed for *d*_5_-IAA.

LC-MS/MS analysis of *d*_5_-IAA was carried out using a system consisting of a quadrupole/time-of-flight tandem mass spectrometer (X500R, AB SCIEX) and an ultrahigh performance liquid chromatography (Nexera, Shimadzu) equipped with a reverse phase column (CORTECS UPLC phenyl, ϕ2.7×100 mm, 1.6 μm, 2.1×75 mm; Waters). For *d*_5_-IAA analysis, the elution of the samples was carried out with H_2_O containing 0.05% AcOH (solvent A) and acetonitrile containing 0.05% AcOH (solvent B), and the mobile phase was changed from 10% B to 40% and 100% at 5 and 7 min after the injection, respectively, at a flow rate of 0.3 mL min^−1^. MS/MS analysis conditions were as follows: Negative ion mode; Declustering potential, −80 V; collision energy, −35 V; and parent ion (*m*/*z*), 174.06 for unlabeled IAA and 179.09 for labeled IAA.

### Quantificational analysis of IAA in *O. minor* seeds treated with auxin biosynthetic inhibitors

*O. minor* seeds were weighed into 1.5 mL micro tube and treated with 200 μL of 0.1% agar. They were covered with aluminum foil and conditioned for 2 weeks at 23°C. Yucasin DF and kynurenine were diluted in 200 nM GR24 (0.1% acetone/water) to 100 μM and 20 μM, respectively. It was added to 200 μL conditioned seeds (final concentration: 100 nM GR24, 50 μM yucasin DF, 10 μM kynurenine), Incubated at 23°C for 5 days. After 5 days, the supernatant was centrifuged and discarded, washed twice with 1 mL sterile water, and then 1 mL acetone was added for overnight extraction at 4°C. After adding 100 pg/mL ^13^*C*_6_-IAA as an internal standard, the supernatant of the acetone extract was collected, concentrated, and dissolved in H_2_O containing 1% AcOH This solution was loaded onto an HLB cartridge column (1 cc, 30 mg, Waters), washed with 1 mL of H_2_O containing 1% AcOH and eluted with 2 mL of 80% MeCN containing 1% AcOH. The eluted fraction was concentrated again and redissolved in MeCN. The sample was subjected to LC-MS/MS to quantify the amount of endogenous IAA. LC-MS/MS analysis of IAA and ^13^*C*_6_-IAA was carried out using the same system that was used for *d*_5_-IAA analysis as mentioned above. MS/MS analysis conditions were as follows: Negative ion mode; Declustering potential, 80 V; collision energy, 15 V; and parent ion (*m/z*), 176.07 for unlabeled IAA and 182.09 for ^13^*C*_6_-IAA.

## Acknowledgments

We thank Dr Steven Runo for kindly providing seeds of *Striga hermonthica*. We thank Dr Huw Tyson from Edanz (https://jp.edanz.com/ac) for editing a draft of this manuscript.

## References

Conn, C.E., Bythell-Douglas, R., Neumann, D., Yoshida, S., Whittington, B., Westwood, J.H., et al. (2015) PLANT EVOLUTION. Convergent evolution of strigolactone perception enabled host detection in parasitic plants. Science 349: 540–543.

de Saint Germain, A., Jacobs, A., Brun, G., Pouvreau, J.B., Braem, L., Cornu, D., et al. (2021) A Phelipanche ramosa KAI2 protein perceives strigolactones and isothiocyanates enzymatically. Plant Commun 2: 100166.

Hayashi, K., Neve, J., Hirose, M., Kuboki, A., Shimada, Y., Kepinski, S., et al. (2012) Rational design of an auxin antagonist of the SCF(TIR1) auxin receptor complex. ACS Chem Biol 7: 590–598.

He, W., Brumos, J., Li, H., Ji, Y., Ke, M., Gong, X., et al. (2011) A small-molecule screen identifies L-kynurenine as a competitive inhibitor of TAA1/TAR activity in ethylene-directed auxin biosynthesis and root growth in Arabidopsis. Plant Cell 23: 3944–3960.

Hylova, A., Pospisil, T., Spichal, L., Mateman, J.J., Blanco-Ania, D. and Zwanenburg, B. (2019) New hybrid type strigolactone mimics derived from plant growth regulator auxin. N Biotechnol 48: 76–82.

Kuruma, M., Suzuki, T. and Seto, Y. (2021) Tryptophan derivatives regulate the seed germination and radicle growth of a root parasitic plant, Orobanche minor. Bioorg Med Chem Lett 43: 128085.

Ogawa, S., Cui, S., White, A.R.F., Nelson, D.C., Yoshida, S. and Shirasu, K. (2022) Strigolactones are chemoattractants for host tropism in Orobanchaceae parasitic plants. Nat Commun 13: 4653.

Steenackers, W., El Houari, I., Baekelandt, A., Witvrouw, K., Dhondt, S., Leroux, O., et al. (2019) cis-Cinnamic acid is a natural plant growth-promoting compound. J Exp Bot 70: 6293–6304.

Steenackers, W., Klima, P., Quareshy, M., Cesarino, I., Kumpf, R.P., Corneillie, S., et al. (2017) cis-Cinnamic Acid Is a Novel, Natural Auxin Efflux Inhibitor That Promotes Lateral Root Formation. Plant Physiol 173: 552–565.

Suzuki, T., Kuruma, M. and Seto, Y. (2022) A New Series of Strigolactone Analogs Derived From Cinnamic Acids as Germination Inducers for Root Parasitic Plants. Front Plant Sci 13: 843362.

Takei, S., Uchiyama, Y., Burger, M., Suzuki, T., Okabe, S., Chory, J., et al. (2023) A Divergent Clade KAI2 Protein in the Root Parasitic Plant Orobanche minor Is a Highly Sensitive Strigolactone Receptor and Is Involved in the Perception of Sesquiterpene Lactones. Plant Cell Physiol.

Toh, S., Holbrook-Smith, D., Stogios, P.J., Onopriyenko, O., Lumba, S., Tsuchiya, Y., et al. (2015) Structure-function analysis identifies highly sensitive strigolactone receptors in Striga. Science 350: 203–207.

Tsuchiya, Y. (2018) Small Molecule Toolbox for Strigolactone Biology. Plant Cell Physiol 59: 1511–1519.

Tsuchiya, Y., Yoshimura, M., Sato, Y., Kuwata, K., Toh, S., Holbrook-Smith, D., et al. (2015) PARASITIC PLANTS. Probing strigolactone receptors in Striga hermonthica with fluorescence. Science 349: 864–868.

Tsuda, E., Yang, H., Nishimura, T., Uehara, Y., Sakai, T., Furutani, M., et al. (2011) Alkoxy-auxins are selective inhibitors of auxin transport mediated by PIN, ABCB, and AUX1 transporters. J Biol Chem 286: 2354–2364.

Tsugafune, S., Mashiguchi, K., Fukui, K., Takebayashi, Y., Nishimura, T., Sakai, T., et al. (2017) Yucasin DF, a potent and persistent inhibitor of auxin biosynthesis in plants. Sci Rep 7: 13992.

Uchida, N., Takahashi, K., Iwasaki, R., Yamada, R., Yoshimura, M., Endo, T.A., et al. (2018) Chemical hijacking of auxin signaling with an engineered auxin-TIR1 pair. Nat Chem Biol 14: 299–305.

Uraguchi, D., Kuwata, K., Hijikata, Y., Yamaguchi, R., Imaizumi, H., Am, S., et al. (2018) A femtomolar-range suicide germination stimulant for the parasitic plant Striga hermonthica. Science 362: 1301–1305.

Wang, D., Pang, Z., Yu, H., Thiombiano, B., Walmsley, A., Yu, S., et al. (2022) Probing strigolactone perception mechanisms with rationally designed small-molecule agonists stimulating germination of root parasitic weeds. Nat Commun 13: 3987.

Xie, X., Yoneyama, K. and Yoneyama, K. (2010) The strigolactone story. Annu Rev Phytopathol 48: 93–117.

Yamada, R., Murai, K., Uchida, N., Takahashi, K., Iwasaki, R., Tada, Y., et al. (2018) A Super Strong Engineered Auxin-TIR1 Pair. Plant Cell Physiol 59: 1538–1544.

