## supplemental figures for "Radicle growth regulation of root parasitic plants by auxin-related compounds"

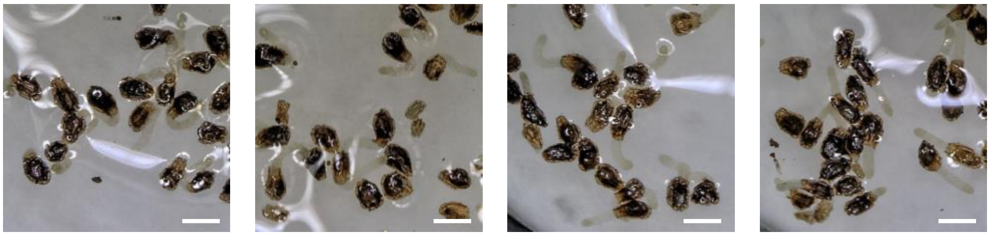

| | | | | |
|---|---|---|---|---|
| *rac*-GR24 (nM) | 10 | 10 | 10 | 10 |
| NPA (μM) | 0 | 0.1 | 1 | 10 |

**Fig. S1**. Effects of the auxin transport inhibitor, NPA, on radicle growth or angle of germinated *O. minor* seeds.. Scale bar = 0.3 mm

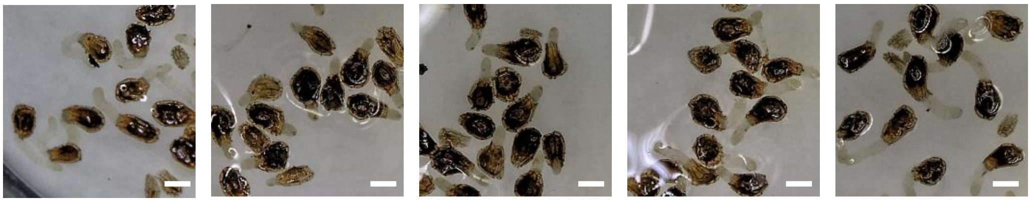

| | | | | | |
|---|---|---|---|---|---|
| *rac*-GR24 (nM) | 10 | 10 | 10 | 10 | 10 |
| IAA (μM) | - | 1 | 1 | 1 | 1 |
| Auxinole (μM) | - | - | 1 | 10 | 100 |

**Fig. S2**. Effects of auxin antagonist on radicle growth of germinated *O. minor* seeds in the presence of exogenously applied IAA. Scale bar = 0.2 mm

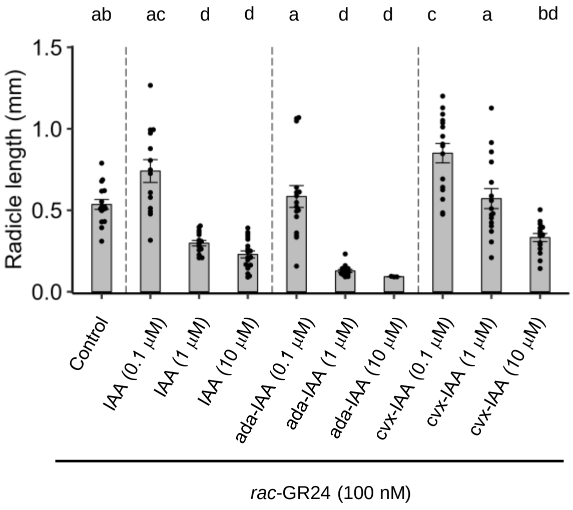


**Fig. S3**. Effects of synthetic auxin analogs on radicle growth of germinated *S. hermonthica*. 100 nM *rac*-GR24 solution contained 0.1% acetone was used as control. Data are the means ± SE ($n$ = 3-19) Different letters indicate significant differences at $P < 0.05$ with Tukey multiple comparison test. Dots show the exact data for individual samples.

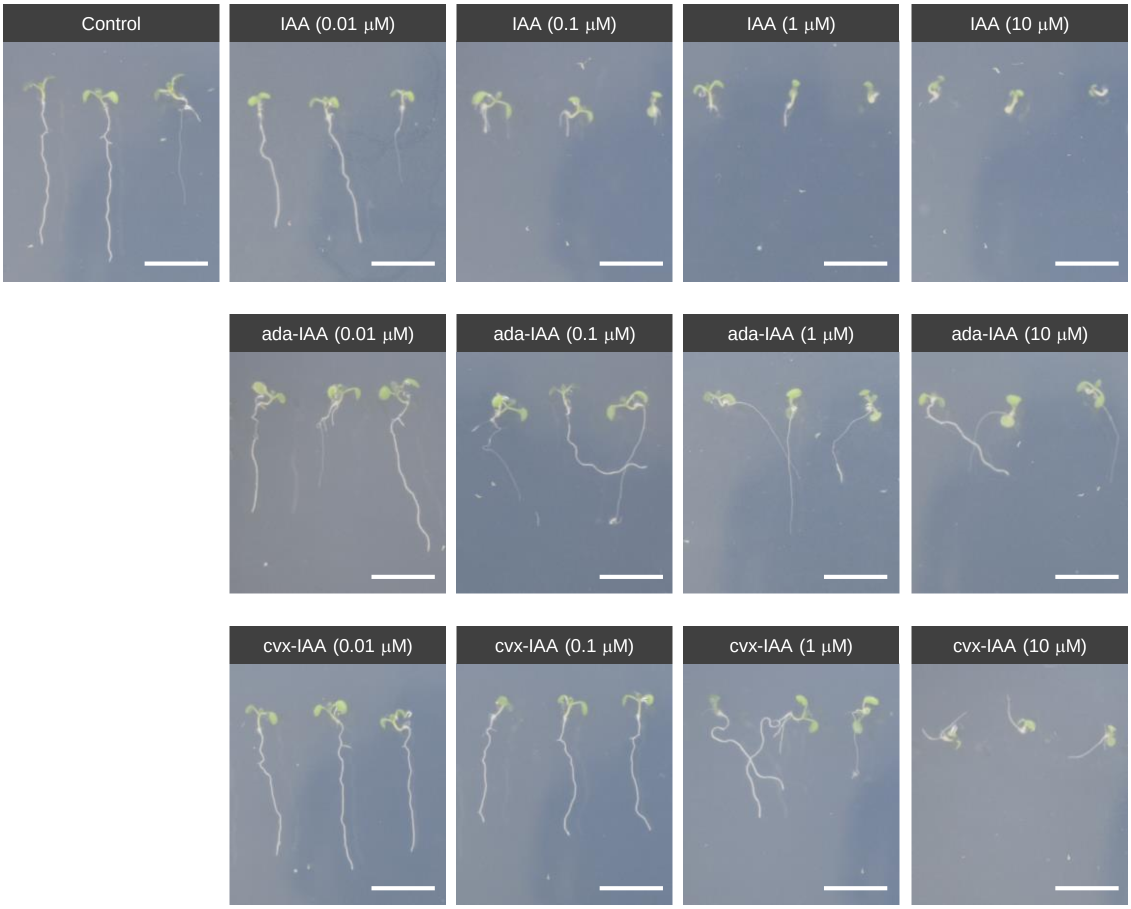


**Fig. S4**. Effects of synthetic auxin analogs on primary root growth of *A. thaliana* seedlings. Scale bar = 10 mm
